# Mutations in nucleoporin *NUP88* cause lethal neuromuscular disorder

**DOI:** 10.1101/347179

**Authors:** Edith Bonnin, Pauline Cabochette, Alessandro Filosa, Ramona Jühlen, Shoko Komatsuzaki, Mohammed Hezwani, Achim Dickmanns, Valérie Martinelli, Marjorie Vermeersch, Lynn Supply, Nuno Martins, Laurence Pirenne, Gianina Ravenscroft, Marcus Lombard, Sarah Port, Christiane Spillner, Sandra Janssens, Ellen Roets, Jo Van Dorpe, Martin Lammens, Ralph H. Kehlenbach, Ralf Ficner, Nigel Laing, Katrin Hoffmann, Benoit Vanhollebeke, Birthe Fahrenkrog

## Abstract

Nucleoporins build the nuclear pore complex (NPC), which, as sole gate for nuclear-cytoplasmic exchange, are of outmost importance for normal cell function. Defects in the process of nucleocytoplasmic transport or in its machinery have been frequently described in human diseases, such as cancer and neurodegenerative disorders, but only in a few cases of developmental disorders. Here we report biallelic mutations in the nucleoporin *NUP88* as a novel cause of lethal fetal akinesia deformation sequence (FADS) in two families. FADS comprises a spectrum of clinically and genetically heterogeneous disorders with congenital malformations related to impaired fetal movement. We show that genetic disruption of *nup88* in zebrafish results in pleiotropic developmental defects reminiscent of those seen in affected human fetuses, including locomotor defects as well as defects at neuromuscular junctions. Phenotypic alterations become visible at distinct developmental stages, both in affected human fetuses and in zebrafish, whereas early stages of development are apparently normal. The zebrafish phenotypes caused by nup88 deficiency are only rescued by expressing wild-type nup88 and not the disease-linked mutant forms of nup88. Furthermore, using human and mouse cell lines as well as immunohistochemistry on fetal muscle tissue, we demonstrate that NUP88 depletion affects rapsyn, a key regulator of the muscle nicotinic acetylcholine receptor at the neuromuscular junction. Together, our studies provide the first characterization of NUP88 in vertebrate development, expand our understanding of the molecular events causing FADS, and suggest that variants in *NUP88* should be investigated in cases of FADS.

## Introduction

The nucleoporin NUP88 [MIM 602552] is a constituent of the nuclear pore complex (NPC), the gate for all trafficking between the nucleus and the cytoplasm (1). NUP88 resides on both the cytoplasmic and the nuclear side of NPCs (2) and it is found in distinct sub-complexes: on the cytoplasmic face it associates with NUP214 [MIM 114350] and NUP62 [MIM 605815] as well as NUP98 [MIM 601021], while on the nuclear side NUP88 binds the intermediate filament protein lamin A [MIM 150330] (2–5). The NUP88-NUP214 complex plays an important role in the nuclear export of a subset of proteins and pre-ribosomes, which is mediated by the nuclear export receptor CRM1 (Required for chromosome maintenance, alias exportin 1, XPO1 [MIM 602559]) (6–8). Depletion of NUP88 alters the intracellular localization of NF-κB proteins (9–11). Moreover, NUP88 is frequently overexpressed in a variety of human cancers and its role therein appears linked to the deregulation of the anaphase promoting complex (12, 13) and its binding to vimentin (14).

Fetal movement is a prerequisite for normal fetal development and growth. Intrauterine movement restrictions cause a broad spectrum of disorders characterized by one or more of the following features: contractures of the major joints (arthrogryposis), pulmonary hypoplasia, facial abnormalities, hydrops fetalis, pterygia, polyhydramnios and *in utero* growth restriction (15). The unifying feature is a reduction or lack of fetal movement, giving rise to the term fetal akinesia deformations sequence (FADS [OMIM 208150]) (16). FADS is a clinically and genetically heterogeneous condition of which the traditionally named Pena-Shokeir subtype is characterized by multiple joint contractures, facial abnormalities, and lung hypoplasia resulting from the decreased *in utero* movement of the fetuses (15). Affected fetuses are often lost as spontaneous abortions (*in utero* fetal demise) or stillborn. Many of those born alive are premature and die shortly after birth. In the past, the genetic basis for these disorders was frequently unknown, but due to the recent availability of next generation sequencing, the molecular etiology is becoming increasingly understood. Many cases of FADS result from impairment along the neuromuscular axis and from mutations in genes encoding components of the motor neurons, peripheral nervous system, neuromuscular junction and the skeletal muscle. Genes encoding components critical to the neuromuscular junction and acetylcholine receptor (AChR) clustering represent a major class of FADS disease genes, these include *RAPSN* [MIM 601592] (17, 18), *DOK7* [MIM 610285] (19), and *MUSK* [MIM 601296] (20), as well as mutations in the subunits of the muscular nicotinic acetylcholine receptor (AChR) (17, 21). These mutations are expected to affect neuromuscular junctions (22).

Here, we report a Mendelian, lethal developmental human disorder caused by mutations in *NUP88*. We demonstrate that biallelic mutations in *NUP88* are associated with fetal akinesia of the Pena-Shokeir-like subtype. We confirm in zebrafish that loss of NUP88 impairs locomotion behavior and that the human mutant alleles are functionally null. We show that loss of NUP88 affects protein levels and localization of rapsyn in cell lines and subject samples. Consistent with altered rapsyn, AChR clustering in zebrafish is abnormal. We propose that defective NUP88 function in FADS impairs neuromuscular junction formation.

## Results

### Identification of NUP88 mutations in individuals affected by fetal akinesia

We performed exome sequencing and Sanger sequencing on genomic DNA from individuals affected with FADS from two families (Figure 1A). Clinical and genetic findings are summarized in Table1, pictures of two fetuses are shown in Figures 1B and 1C, pedigrees and gene structure are shown in Figures 1A and 1D. Family A comprises four affected individuals, three male and one female (Figure 1A; A.II.3, 4, 5, 7), and four healthy siblings born to consanguineous parents of Palestinian origin. Exome sequencing of the last affected fetus A.II.7 revealed a homozygous missense mutation c.1300G>T (p.D434Y) in the *NUP88* gene [NM_002532.5] (Figure 1A), absent in relevant databases (dbSNP, Ensembl, UCSC, TGP, ExAC, HGMD, gnomAD). Sanger sequencing revealed identical homozygous missense mutation in the third affected fetus (Figure 1A, A.II.5; Figure S1A). Both parents and unaffected siblings A.II.1, A.II.2 and A.II. 6 are heterozygous carriers of the mutation, unaffected sibling A.II.8 carries two intact alleles of *NUP88* after *in vitro* fertilization and preimplantation diagnostic (Figure 1A). DNA was unavailable from the first and second miscarriage (A.II.3 and A.II.4), but clinical phenotypes resemble those of the two affected individuals A.II.5 and A.II.7 (Table 1). In Family B, one affected son was born to healthy unrelated parents of European descent. Exome sequencing in the affected individual, his parents and his two unaffected sibs (Figure S1B) revealed that the individual is compound heterozygous for two *NUP88* mutations, i.e. a nonsense c.1525C>T (p.R509*) and a frameshift c.1899-1901del (p.E634del; Figure 1A; B.II.2), originally absent in relevant databases. Variants were entered in dbSNP and gnomAD. Parents and healthy siblings were heterozygous carriers of the one or the other of the mutations, thus confirming correct segregation consistent with recessive inheritance (Figure 1A).

**Table 1:**
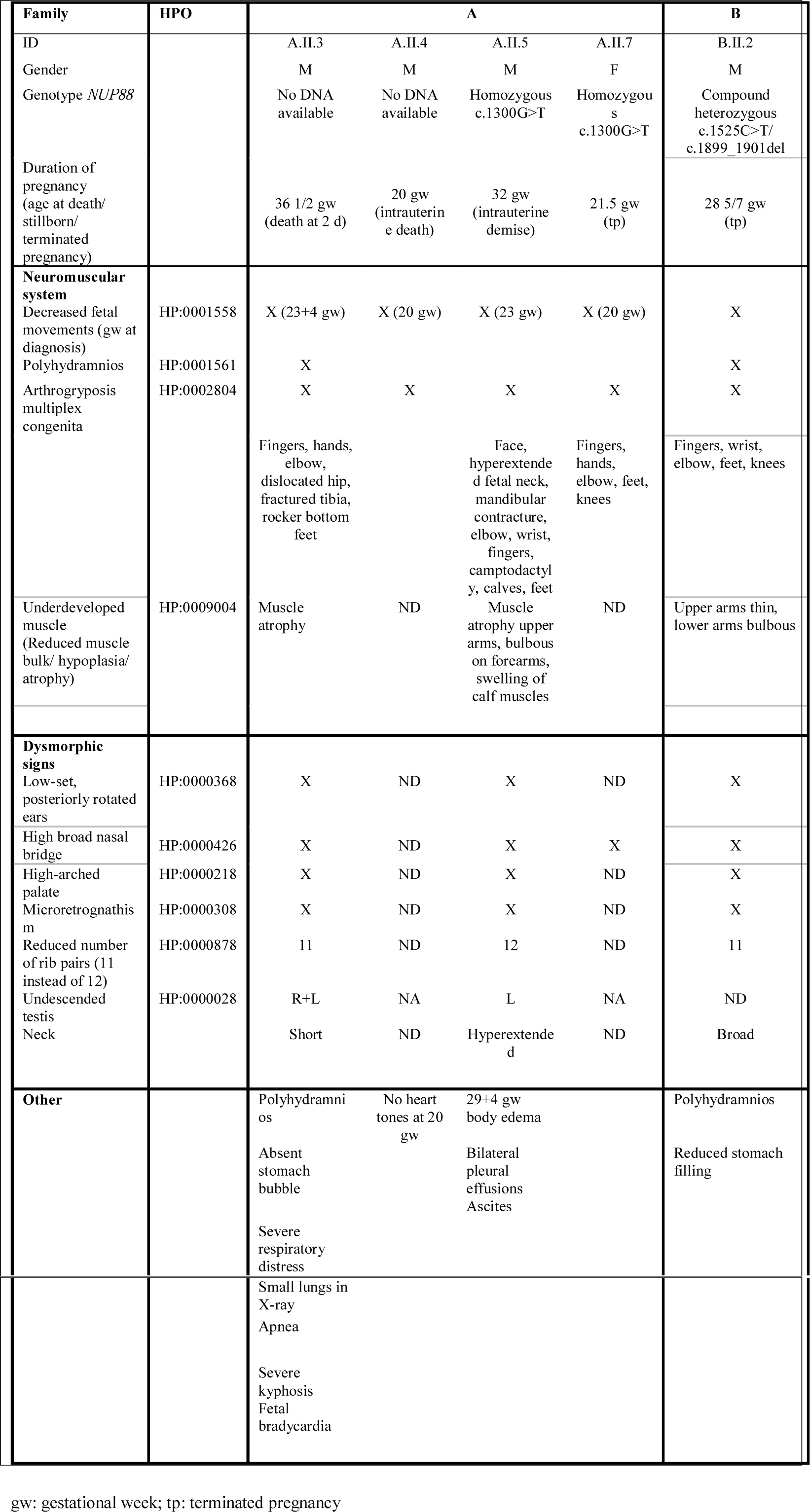
Genetic and Clinical Data of Affected Individuals with *NUP88* Variants

| Family | HPO | A |  |  |  | B |
| --- | --- | --- | --- | --- | --- | --- |
| ID |  | A.II.3 | A.II.4 | A.II.5 | A.II.7 | B.II.2 |
| Gender |  | M | M | M | F | M |
| Genotype *NUP88* |  | No DNA available | No DNA available | Homozygous c.1300G>T | Homozygous c.1300G>T | Compound heterozygous c.1525C>T/ c.1899_1901del |
| Duration of pregnancy (age at death/ stillborn/ terminated pregnancy) |  | 36 1/2 gw (death at 2 d) | 20 gw (intrauterine death) | 32 gw (intrauterine demise) | 21.5 gw (tp) | 28 5/7 gw (tp) |
| **Neuromuscular system** |  |  |  |  |  |  |
| Decreased fetal movements (gw at diagnosis) | HP:0001558 | X (23+4 gw) | X (20 gw) | X (23 gw) | X (20 gw) | X |
| Polyhydramnios | HP:0001561 | X |  |  |  | X |
| Arthrogryposis multiplex congenita | HP:0002804 | X | X | X | X | X |
|  |  | Fingers, hands, elbow, dislocated hip, fractured tibia, rocker bottom feet |  | Face, hyperextended fetal neck, mandibular contracture, elbow, wrist, fingers, camptodactyly, calves, feet | Fingers, hands, elbow, feet, knees | Fingers, wrist, elbow, feet, knees |
| Underdeveloped muscle (Reduced muscle bulk/ hypoplasia/ atrophy) | HP:0009004 | Muscle atrophy | ND | Muscle atrophy upper arms, bulbous on forearms, swelling of calf muscles | ND | Upper arms thin, lower arms bulbous |
| **Dysmorphic signs** |  |  |  |  |  |  |
| Low-set, posteriorly rotated ears | HP:0000368 | X | ND | X | ND | X |
| High broad nasal bridge | HP:0000426 | X | ND | X | X | X |
| High-arched palate | HP:0000218 | X | ND | X | ND | X |
| Microretrognathism | HP:0000308 | X | ND | X | ND | X |
| Reduced number of rib pairs (11 instead of 12) | HP:0000878 | 11 | ND | 12 | ND | 11 |
| Undescended testis | HP:0000028 | R+L | NA | L | NA | ND |
| Neck |  | Short | ND | Hyperextended | ND | Broad |
| **Other** |  | Polyhydramnios | No heart tones at 20 gw | 29+4 gw body edema |  | Polyhydramnios |
|  |  | Absent stomach bubble |  | Bilateral pleural effusions |  | Reduced stomach filling |
|  |  | Severe respiratory distress |  | Ascites |  |  |

|  |  |  |
| --- | --- | --- |
|  |  | Small lungs in<br>X-ray<br>Apnea<br>Severe<br>kyphosis<br>Fetal<br>bradycardia |

gw: gestational week; tp: terminated pregnancy

**Figure 1:**
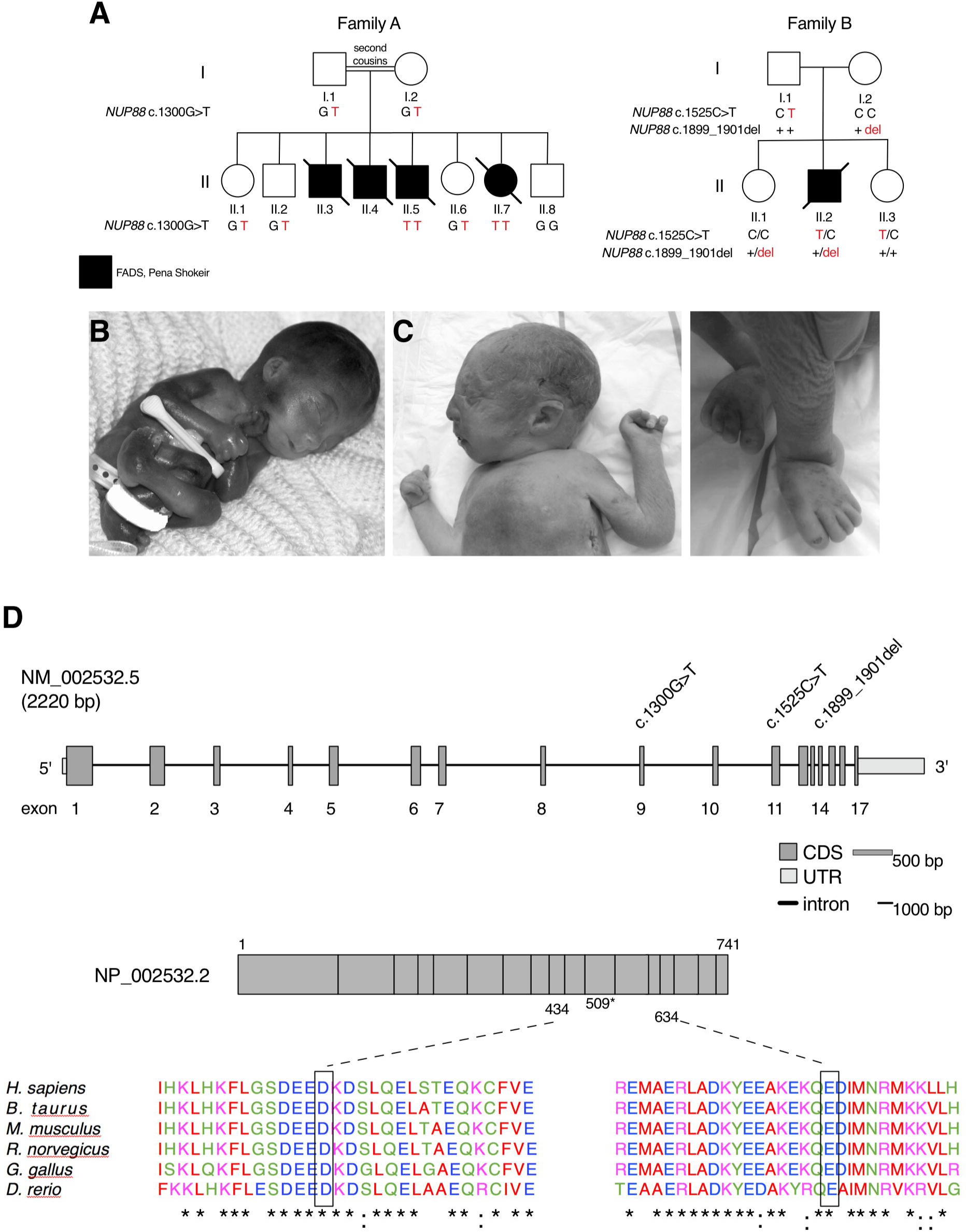
*NUP88* mutations identified in affected individuals from two families. (**A**) Pedigrees of two families identified with mutations in *NUP88* (GenBank: NM_002532.5). Macroscopic appearance of (**B**) one fetus (II.7) homozygous for the c.1300G>T variant and (**C**) the heterozygous fetus harboring the c.1525C>T and c.1899_1901del variants in *NUP88*. Note the multiple contractures and facial anomalies. (**D**) *NUP88* gene and protein structure, location of the identified mutations, and phylogenetic conservation of the mutated residues and surrounding amino acids. Identical amino acids are indicated by asterisks, highly similar residues by colons.

### Structural modelling of NUP88

The missense substitution p.D434Y and deletion p.E634del affect evolutionary highly conserved NUP88 residues (Figure 1D) indicating functional relevance. Accordingly, SIFT/Provean, Polyphen-2, and MutationTaster predicted both mutations to be disease causing or potentially pathogenic (Table S1). The crystal structure of NUP88 is not known, therefore we performed structural modelling. Models obtained (see Methods) predicted the N-terminal domain (NTD) to form a 7-bladed ß-propeller, set up in a (4, 4, 4, 4, 4, 4, 3) arrangement of ß-strands and no Velcro lock as typical for classical ß-propellers (Figure 2A). Around 60 residues precede the ß-propeller and are located at the bottom or side of the propeller thereby shielding 2-4 blades in their vicinity (Figure 2A). The model reveals high similarity to the PDB deposited structures of Nup82 from Baker’s yeast and Nup57 from *Chaetomium thermophilum* (Figure 2B). The most prominent differences are a loop region and a helix-turn-helix (HTH) motif emanating from blades 4 and 5, respectively (Figure 2B). Models obtained for NUP88’s C-terminal domain (CTD) exhibited low reliability, but the CTD, in analogy to its yeast homolog, is likely composed of extended α-helices (Figure 2C) that form trimeric coiled-coils, either in *cis* or in *trans*. In this context, an arrangement with its complex partners NUP214 and NUP62 in *trans* is most likely, as described for the yeast counterpart of the complex (23, 24).

**Figure 2:**
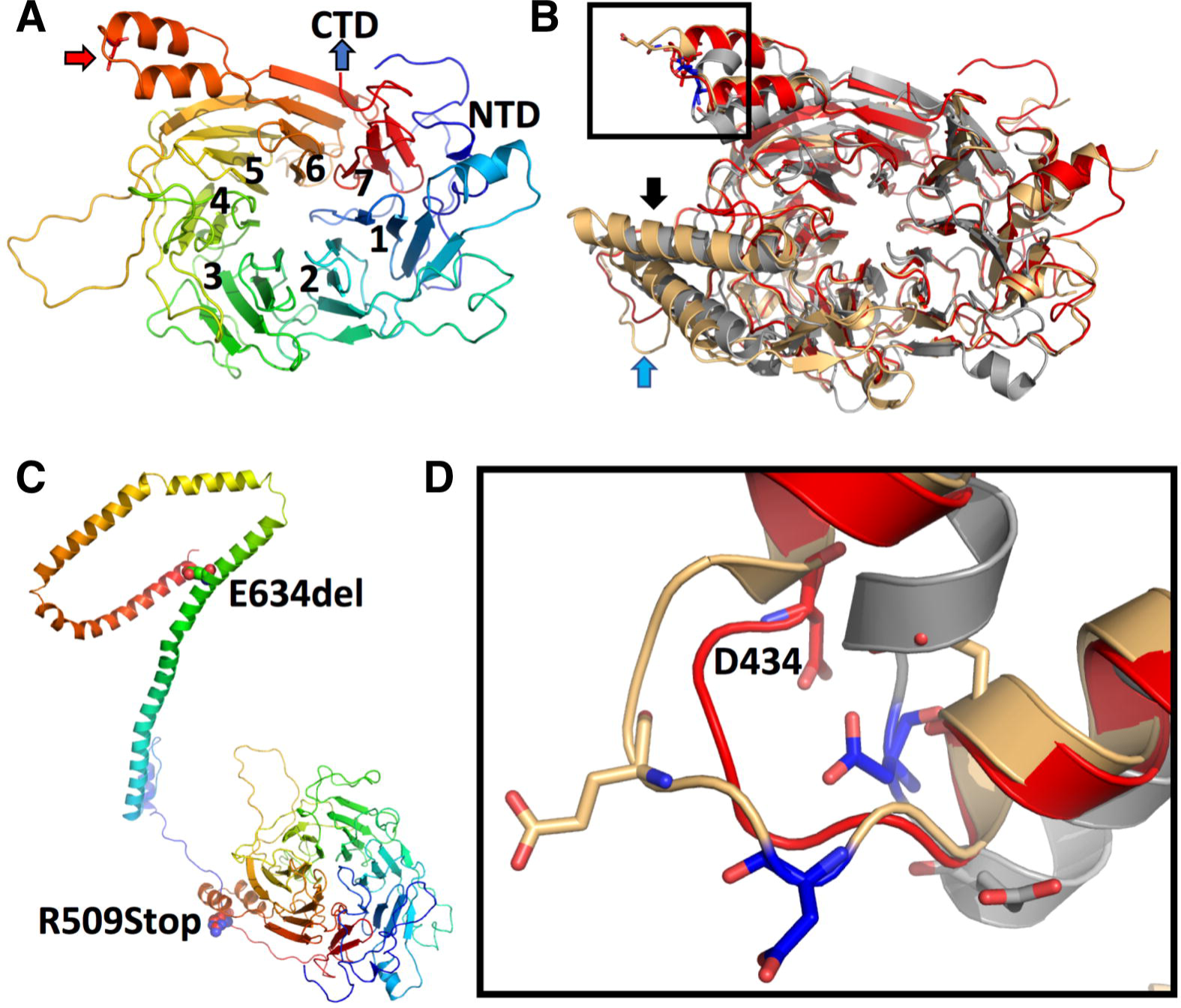
Modelling of human NUP88. (**A**) The N-terminal domain of NUP88 reveals a seven-bladed ß-propeller with an N-terminal extension. The rainbow coloring indicates N-terminal residues in blue (NTD = N-terminal domain) and C-terminal residues of the propeller in red. Individual blades are indicated by numbers. The red arrow indicates the location of the p.D434Y point mutation (see below for details). (**B**) Overlay of the NUP88 model (red) with the X-ray structures of Nup82 from baker’s yeast (PDBid: 3pbp, in gray) and *C. thermophiles* (PDBid: 5cww; in light yellow). Significant differences between species are in blade 4 (HTH-motif; black arrow) and 5 (extended loop; blue arrow). (**C**) Composite model of the N- and C-terminal regions of NUP88. The presented model was generated using RaptorX with its standard settings and misses about 40 amino acid residues after the propeller region. Both, the propeller and CTD regions are colored in rainbow coloring as in (**A**). The individual mutations are indicated by their numbering and represented in sphere mode. (**D**) Magnification of the loop bearing the D434 mutation in NUP88 in stick mode. The coloring of the individual molecules is as described in (**B**).

According to the model structure, the p.D434Y mutation is located in the loop of a HTH motif between the two outermost ß-strands of blade 6 (Figure 2B, overall view; Figure 2D, magnification). The mutation likely leads to a decrease in the interaction with one of the neighboring proteins, thereby leading to a destabilization of the complex. The non-sense mutation c.1525C>T resulting in p.R509* is located just after the ß-propeller in the linker region to the CTD resulting in a complete loss of all α-helices. Thus, the interaction of NUP88 with its complex partners is likely reduced to only propeller interactions, if the protein is not completely lost due to non-sense mediated decay of the mRNA. The p.E634del mutation is located in the middle of the CTD sequence and predicted to lie in the last fifth of an extended helix. The deletion results in a frame-shift of the remainder of the α-helix, which shifts the following residues by about a third of a helical turn and thus disrupts the interaction pattern of all following residues, which, as a consequence, decreases the overall stability of the interactions within this helix bundle.

### Analysis of nup88 expression during Danio rerio development

To study the function of NUP88 in vertebrate development, we used a zebrafish (*D. rerio*) model. The single zebrafish *nup88* orthologue (ENSDARG00000003235) encodes a protein of 720 amino acid translated from a single 2410 bp transcript. The predicted translated gene product shares 63% identity and 75% similarity with human NUP88. Whole-mount *in situ* hybridization (WISH) and RT-PCR analysis in wild-type AB zebrafish showed that *nup88* transcripts are maternally inherited (Figures S2A and S2B, four-cell-stage embryos) and ubiquitously expressed at 5 hours post fertilization (hpf). By 24 hpf, high levels of *nup88* mRNA were detected in highly proliferative frontal regions of the embryo, i.e. the central nervous system, brain, eye and anterior trunk. At 72 hpf, *nup88* transcript levels are decreasing in these frontal regions. Similar expression patterns in the developing zebrafish have been described for the two NUP88-binding partner, NUP98 and NUP62 (25, 26).

### Genetic disruption of nup88 affects zebrafish development

To study the impact of *nup88* deletion on zebrafish development, we used the *nup88^sa2206^* allele generated by the Zebrafish Mutation Project (27, 28). Heterozygous *nup88^sa2206^* carriers were outcrossed for four generations with wild-type AB zebrafish prior to phenotypic analysis. The *nup88^sa2206^* allele is characterized by a nonsense mutation, c.732T>A (Figure 3A), resulting in a premature stop codon at amino acid 244. *nup88* mRNA levels are reduced by about 90% in *nup88* mutants (see Figure 7E), suggesting that the mRNA is subjected to nonsense-mediated decay. For the purpose of this study, *nup88^sa2206/sa2206^* is therefore referred to as *nup88^−/−^*. During early stages of development and up to 3 days post fertilization (dpf), no marked differences in morphological features of *nup88^−/−^* compared to *nup88^+/+^* and *nup88*^+/−^ siblings were observed. Starting at 4 dpf, phenotypic alterations became visible: smaller head and eyes, lack of a protruding mouth, downwards curvature of the anterior-posterior axis, abnormal gut and aplastic swim-bladder (Figure 3B). Further analyses of the cranial abnormalities revealed that *nup88^−/−^* larvae exhibit severe defects in the ventral viscerocranium formed by seven cartilaginous pharyngeal arches (29, 30). In *nup88^−/−^* larvae, the posterior pharyngeal arches 3-7 were dramatically reduced, distorted or even absent (Figure 3C). The reduced size of head and eyes likely correlated with an increase in apoptosis in the head of *nup88^−/−^* embryos (Figure 3D). Apoptotic cells, as assessed by acridine orange staining, were readily detected in the eyes, the brain and the anterior trunk of 35 hpf mutant embryos, but not in other parts of the body (Figure S2C). Together, these data indicate that *nup88* mutants are phenotypically similar to the large class of jaw and branchial arch zebrafish mutants, designated the flathead group (31, 32). Disruption of *nup88* furthermore led to impaired survival with most death of *nup88^−/−^* larvae occurring at or after 9 dpf (Figure 3E).

**Figure 3:**
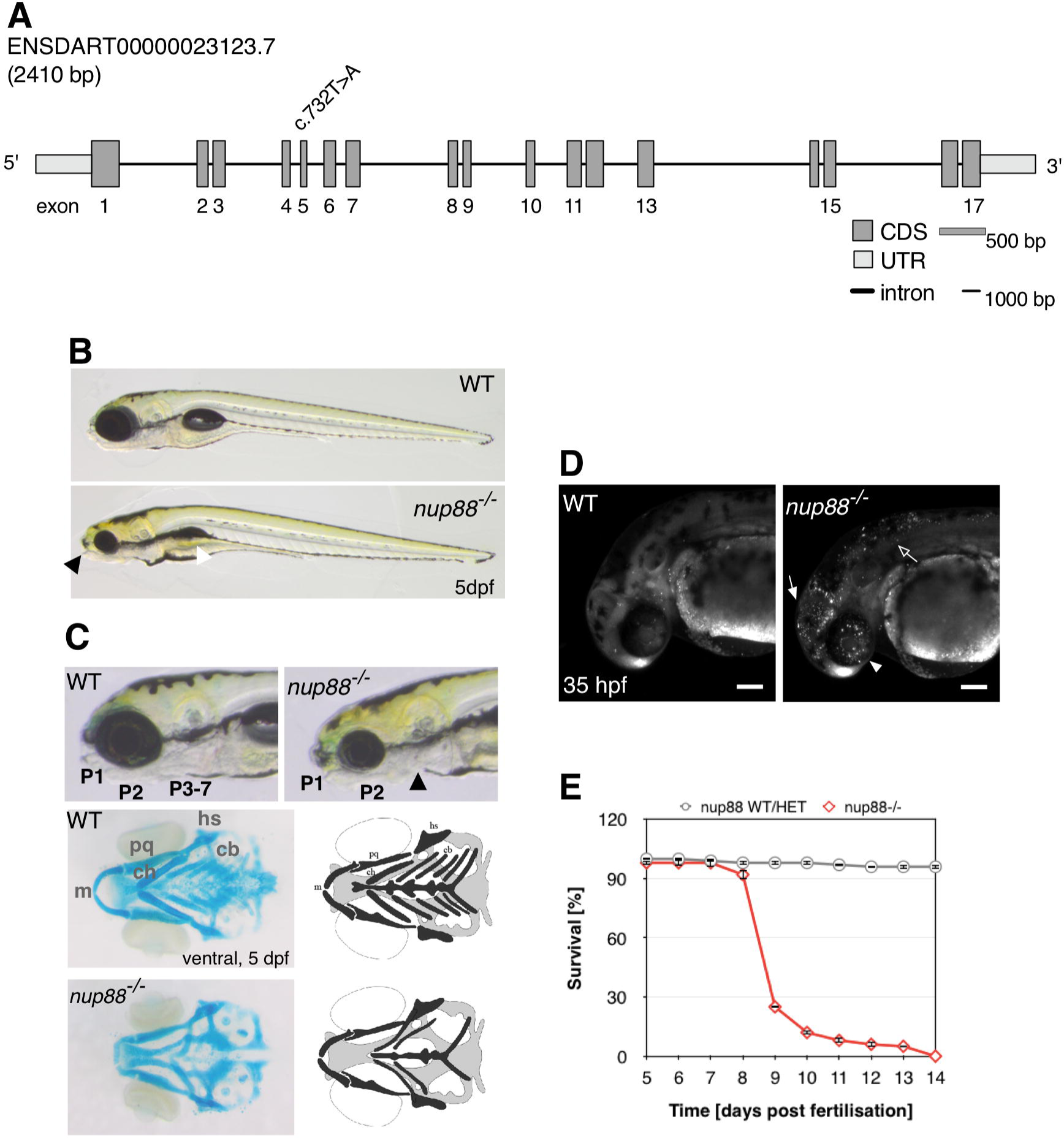
Morphological phenotypes of *nup88^−/−^* mutants. (**A**) *nup88* gene structure and location of the mutation in the *sa2206* allele. (**B**) Lateral view of wild-type and *nup88^−/−^* embryos at 5 dpf. *nup88^−/−^* mutants belong to the flathead group of mutants characterized by decreased head and eye size and the absence of a protruding mouth (black arrowhead). The larvae furthermore show aplastic swim bladder (white arrowhead), hypoplastic liver, abnormal gut and a marked curvature of the anterio-posterior axis. (**C**) Higher magnification lateral views of the head region of wild-type and *nup88^−/−^* embryos at 5 dpf. Alcian blue staining of the viscerocranium revealed that *nup88^−/−^* mutants lack pharyngeal arches 3 to 7 (P3-7). A ventral view of the head showed that hyoid and mandibular arches (P1 and P2) were present, but dysmorphic. A schematic representation of the viscerocranium is shown to illustrate alcian-blue images. Genotypes of fishes were determined by fin clip and RFLP. m, Meckel’s cartilage; ch, ceratohyal; pq, palatoquadrate; cb, ceratobranchials (P 3-7); hs, hyosymplectic. (**D**) Acridine orange staining revealed an increase in apoptotic cells in the head, including the eyes (arrowhead), the brain (filled arrow), and anterior part of the trunk (arrow) of *nup88^−/−^* mutants at 35 hpf compared with wild-type siblings. Shown are confocal images. Scale bars, 100 µm. (**E**) Survival curves of *nup88* mutants and siblings. 75 larvae were analyzed each. Error bars are ±SEM.

### nup88 mutants show strongly impaired locomotor behavior

To further investigate the implication of NUP88 in the etiology of FADS, we next determined whether locomotor function was impaired in *nup88^−/−^* zebrafish using locomotion and touch-evoked escape assays. Zebrafish embryos develop spontaneous muscle contractions at 18 hpf (33), therefore we first analyzed the coiling behavior of *nup88^−/−^* embryos as compared to *nup88*^+/+^ and *nup88*^+/−^ embryos at 22-24 hpf. We did not detect problems in coiling behavior in *nup88^−/−^* embryos at this developmental stage (Figure S3A, Supplementary Movie 1). Next, we analyzed spontaneous swimming activity at 4 dpf (Figure 4A) and found that only about 35% of the *nup88^−/−^* larvae showed spontaneous movement as compared to ~83% of *nup88^+/+^* and about 73% of *nup88^+/−^* larvae (Figure 4B). Moreover, those *nup88^−/−^* larvae moving displayed drastically reduced motor activity, traveled shorter distance (Figure 4C) and initiated less often swim bouts (Figure 4D). In contrast, statistically significant differences in the mean velocity were not observed in *nup88* mutant larvae (Figure 4E).

**Figure 4:**
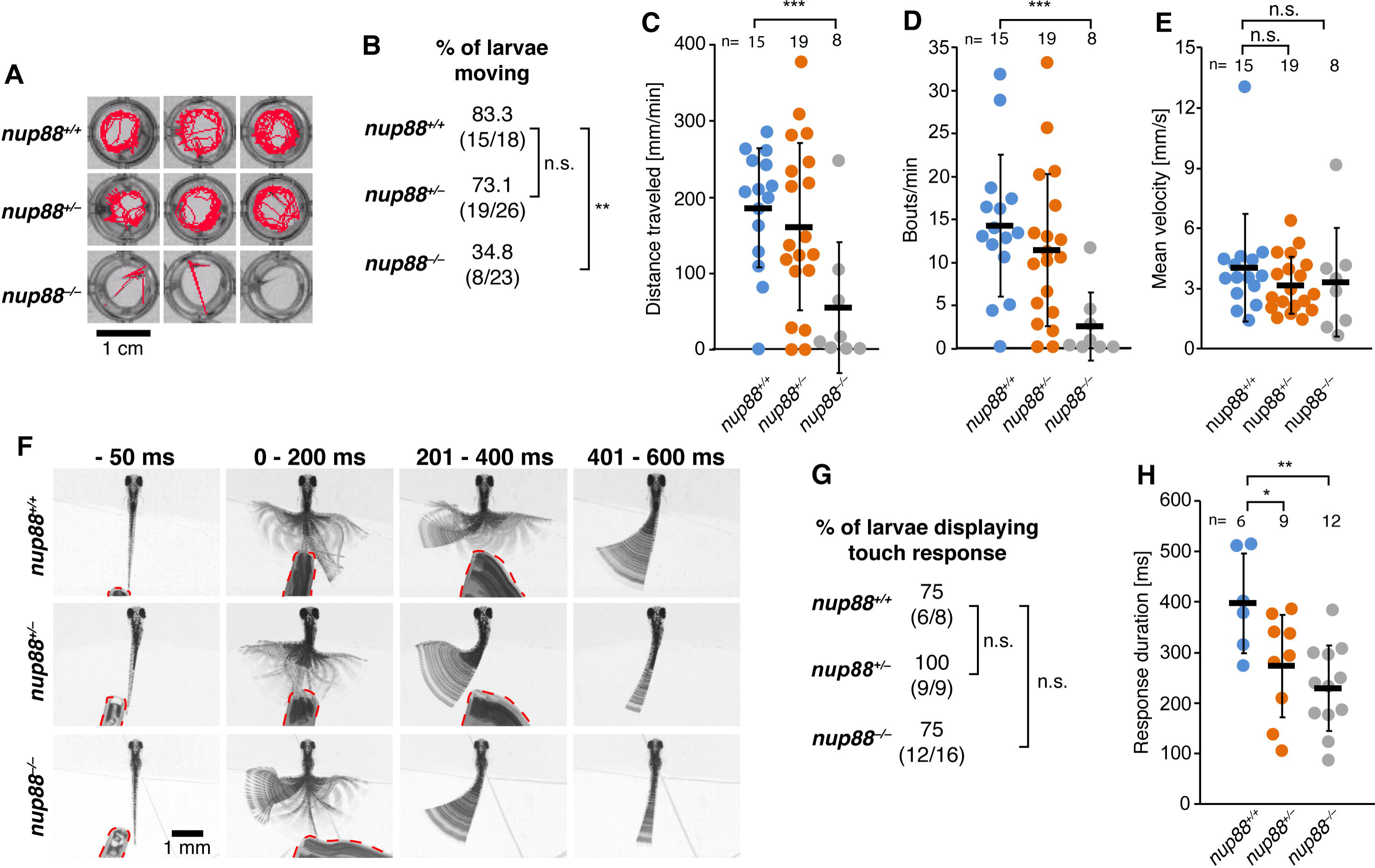
*nup88* mutants display motor impairments. (**A**) Images showing representative examples of motion tracking (red lines) of 4 dpf *nup88* homozygous and heterozygous mutants, and wild-type controls during one-minute long spontaneous locomotion recordings (see also Supplementary Movie 1). (**B**) Quantification of percentages of larvae displaying spontaneous movement, also shown as (number of moving larvae/total number of larvae). Column graphs with individual data points depicting distance travelled/minute (**C**), bouts/minute (**D**), and mean velocity (**E**) of *nup88* mutants and wild-type controls displaying spontaneous locomotion. (**F**) Representative examples of touch-induced escape behavior of 4 dpf head-restrained *nup88* mutant and wild-type larvae. The first column depicts single frames taken 50 ms before an escape response was induced by touching the trunk of larvae with a pipette tip (red dashed lines). The other columns show superimposed frames of escape responses at consecutive time intervals after touch. The asymmetric tail movement displayed in some of the images (*nup88^+/−^* and *nup88^−/−^* at 201-400 ms, and all genotypes at 401-600 ms) represents slow motion of the tail toward its original position. The escape response terminated before this phase (see also Supplementary Movies 2-4). (**G**) Quantification of percentages of larvae displaying touch-induced escape response, also shown as (number of responsive larvae/total number of larvae). (**H)** Graph showing average durations of escape response in *nup88* mutants and wild-types. * p = 0.05, ** p < 0.004, *** p = 0.001, n.s. = not significant, two-tailed Fisher exact test (**B** and **G**) or two-tailed *t*-test (**C-E, H**). Data in **C-E** and **H** are shown as mean ± SD. n is number of embryos/larvae analyzed.

We next performed touch-evoked escape response assays at 3 dpf and 4 dpf. At 3 dpf, percentages of responsive animals and response duration were not significant different between wild-type and *nup88* mutant larvae (Figure S3B). At 4 dpf percentages of responsive animals were also not significant different between wild-type and *nup88* mutant larvae (Figure 4G), however, the response duration among the larvae that moved was significantly reduced in homozygous *nup88* mutants in comparison to wild-type zebrafish. Interestingly also heterozygous *nup88* mutants showed a shortened response duration, although less significant (Figures 4F, 4H; Supplementary Movies 2-4).

### FADS-related mutations in nup88 lead to a loss-of-function phenotype

To address the question whether *NUP88* mutations identified in the familial cases of FADS affect NUP88 function, we performed phenotypic rescue experiments in zebrafish. Two of the three mutated residues in the uncovered FADS cases are conserved between human and zebrafish (Figure 1D), hence we introduced the corresponding mutation on zebrafish expression constructs by site-directed mutagenesis. Human c.1300G>T, p.D434Y corresponds to c.1240G>T, p.D414Y in zebrafish and human c.1899_1901del, p.E634del to zebrafish c.1837_1839del, p.E613del. Human p.R509 is not conserved in zebrafish, therefore we inserted a stop codon at c.1468-1470>TGA, p.H490*, a residue in a similar position as human R509. Subsequently synthetic mRNA corresponding to each variant was microinjected into one-cell stage *nup88*^−/−^ mutants and their rescue capability was assessed by evaluating the eye size as well as the number and morphology of pharyngeal arches. Injection of wild-type (WT) *nup88* mRNA largely rescued the developmental defects of the 5 dpf mutant larvae as indicated by significant restoration of the eye size (Figures 5A and 5B) and a significant increase in the number of pharyngeal arches (Figures 5A and 5C). In addition, the arches largely resembled the morphologically wild-type structures (Figures 5A and 5D). In contrast to WT *nup88* mRNA, injection of *nup88* D414Y mRNA, *nup88* H490* mRNA as well as *nup88* E613del mRNA failed to suppress the *nup88^−/−^* phenotype (Figures 5A-5D), indicating that the resulting nup88 mutant proteins are functionally null.

**Figure 5:**
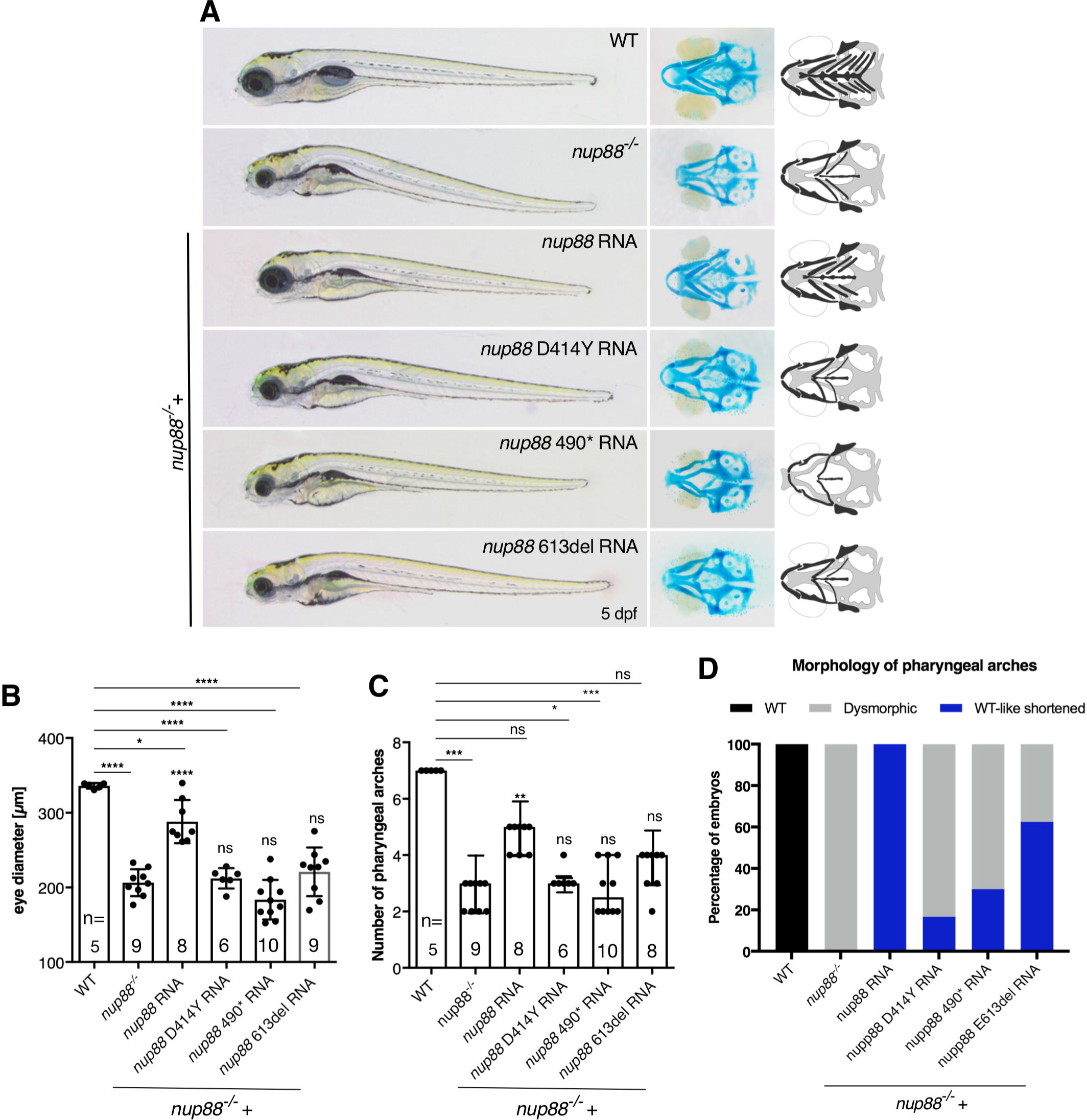
Wild-type *nup88*, but not disease-related mutant forms, rescue defects of *nup88^−/−^* embryos. (**A**) *nup88^+/−^* embryos were in-crossed and 300 pg of mRNA encoding wild-type or the respective mutant *nup88* were microinjected at the one-cell stage. The extent of rescue of each variant mRNA was evaluated after 5 dpf using (**B**) the diameter of the eye, the number of pharyngeal arches (**A** and **C**) and (**D**) their morphology as readouts. Only injecting wild-type, but not the mutant forms of *nup88* mRNA, rescued the reduced eye size and the number of pharyngeal arches as revealed by Alcian blue staining (**A**). At least three independent injections were performed for each condition. Values are mean ± SD. Significance in comparison to *nup88^+/+^* and *nup88^−/−^* embryos, respectively. *P<0.05, **P<0.01, ***P<0.001, ****P<0.0001 (One-way ANOVA test). n is number of embryos/larvae analyzed. In (**D**) 21 larvae from 3 independent experiments (3×7) were analyzed for each form.

### FADS-related mutations in NUP88 alter NUP88 interactions

Consistent with data from higher vertebrates, NUP88 is not essential for NPC integrity: NPCs remain unaffected by the loss of *nup88* in *D. rerio* brain sections (Figure S4A) and in muscle histology sections of individual B.II.2 (Figure S4B) as revealed by immunolabelling with a monoclonal antibody recognizing NPCs (mAb414). To assess whether the *NUP88* mutations identified in the individuals with FADS affect the recruitment of NUP88 to NPCs, we performed immunofluorescence microscopy of GFP-tagged NUP88 protein. Upon expression in HeLa cells, wild-type NUP88 and all mutants were co-localizing with the NPC marker mAb414, although recruitment of the NUP88 p.R509* and p.E634del mutants to NPCs appeared reduced compared to wild-type NUP88 and the p.D434Y mutant (Figure S4C). Moreover, all forms of NUP88 also localized partially to the cytoplasm, as previously seen for NUP88 overexpression (2, 12), and NUP88 p.R509* to the nucleoplasm (Figure S4C). To define the effect of the mutations in NUP88 on the interface with its binding partners NUP214, NUP98 and NUP62, we employed GFP trap affinity purification in combination with Western blot analysis of lysates from HeLa cells expressing the GFP-NUP88 mutants. We found that NUP88 and the p.D434Y mutant co-purified NUP214 and NUP62, while the p.R509* and the p.E634del mutant did not so (Figure 6A). Binding of NUP214 to GFP alone was similar as compared to the p.R509* and the p.E634del mutant, indicating some unspecific binding of the NUP214 and/or the antibodies to GFP. The disrupted interaction between NUP88 and NUP214, however, did not impair NUP214 localization at NPCs (Figure 6B), whereas NUP62 association with NPCs was reduced in cells expressing NUP88 E634del (Figure 6C). Our GFP trap assays further showed that NUP98 associated with NUP88 and all mutant forms (Figure 6A). Consequently, NUP98 association with NPCs appeared unaffected in cells expressing NUP88 mutants (Figure 6D). Furthermore, the disease-related mutations in *NUP88* did not affect the organization of the nuclear lamina as revealed by immunofluorescence analysis of lamin A/C (LA/C; Figure S4D), Western blots analysis of protein levels of lamin A/C and GST-pull-down down assays with NUP88 and the p.D434Y mutant (Figures S4E and S4F). The integrity of the nuclear envelope (NE) was further evidenced by immunofluorescence analyses of the inner nuclear membrane marker Sun1 and Sun2 as well as the outer nuclear membrane marker proteins Nesprin 1 and Nesprin 2 (not shown): the organization of these NE proteins is indistinguishable between HeLa cells overexpressing wild-type NUP88 or any of the disease-related mutants. Depletion of NUP88 from HeLa cells using siRNAs had no visible effect on the distribution of lamin A/C (LA/C) and the NE marker proteins emerin, Nesprin-1, Nesprin-2, Sun1, and Sun2 (Figure S5).

**Figure 6:**
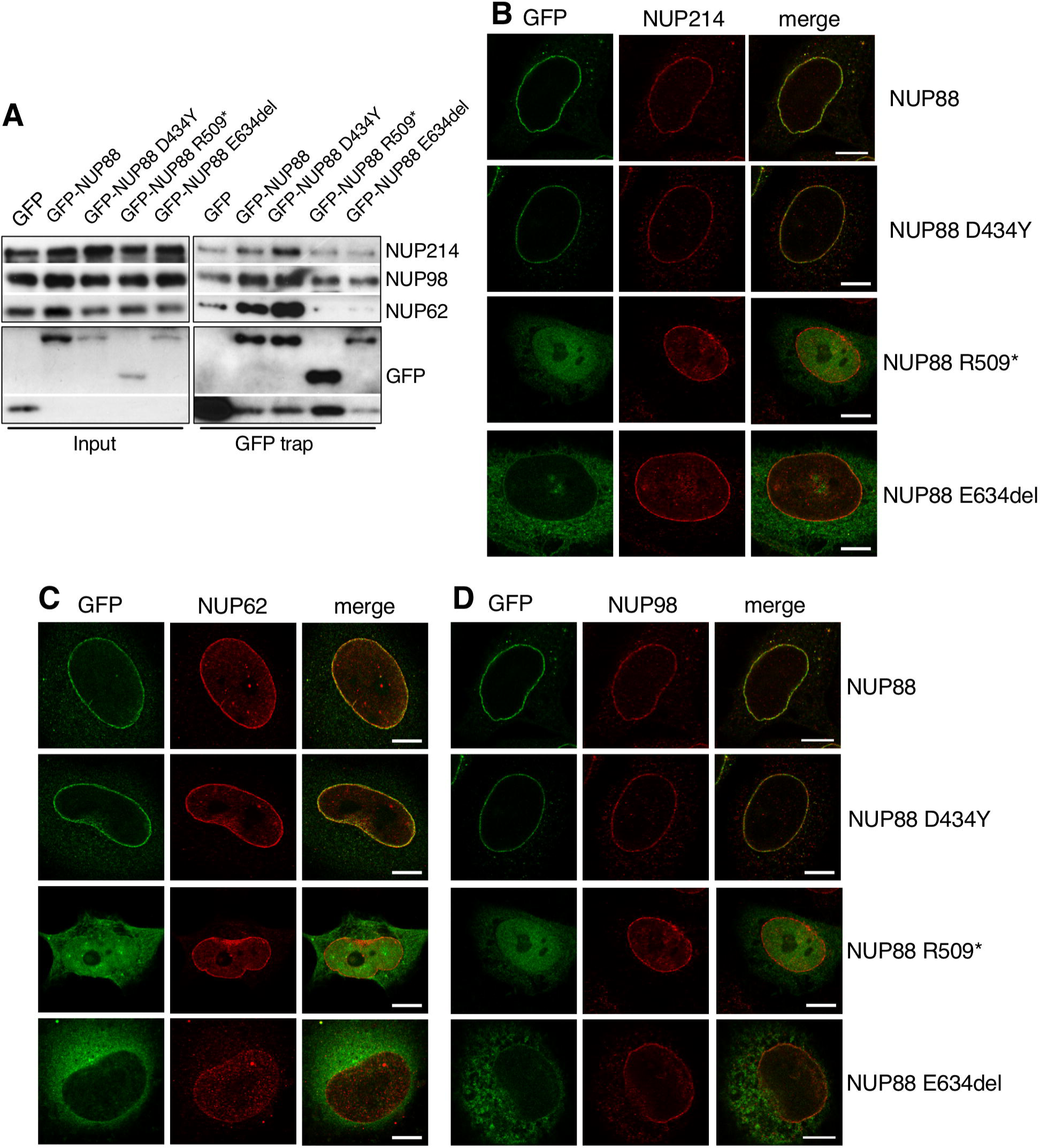
NUP88 mutants have distinct effects on NUP88 nucleoporin partners. (**A**) GFP-trap assays to study the effects of NUP88 mutants on its interaction with NUP214, NUP62 and NUP98. GFP and GFP-NUP88 fusion proteins were transiently expressed in HeLa cells. After 48 h, GFP proteins and associated factors were recovered from cell lysates and probed by Western analysis using antibodies against Nup214, NUP62, and NUP98. Successful transfection and expression of the proteins was confirmed by probing with antibodies against GFP. GFP proteins had the following size: GFP-NUP88 WT, D434Y, and E634del: ~115 (88 + 27) kDa; GFP-NUP88 R509*: 83 (56 + 27) kDa; GFP: 27 kDa. (**B-D**) HeLa cells were transiently transfected with GFP-constructs of the respective NUP88 mutant and fixed and stained after 48 hours for immunofluorescence microscopy using (**B**) anti-NUP214, (**C**) anti-NUP62, and (**D**) anti-NUP98 antibodies. Only NUP88 E634del affects NUP62 association with NPCs. Scale bars: 10 µm

As NUP88 is critically involved in CRM1-dependent nuclear export of proteins, we asked next whether the mutations in NUP88 affect nuclear import and/or export, but we observed no defects in general nuclear protein import or export (Figure S6A) or the three CRM1 targets mTOR, p62/SQSTM and TFEB (Figure S6B).

### Loss of NUP88 affects rapsyn levels and localization

Impeded formation of AChR clusters at the neuromuscular junction (NMJ) is considered a key defect in FADS. Given the central role of rapsyn in AChR clustering and FADS (17, 18, 21), we therefore asked whether a loss of NUP88 function would negatively affect rapsyn and depleted NUP88 by siRNAs from HeLa and C2C12 cells and monitored protein levels of rapsyn by Western blotting. As shown in Figure 7A, depletion of NUP88 from HeLa cells in fact coincided with a decrease in rapsyn levels. Quantification revealed that siRNA treatment led to reduction of NUP88 by 80-90% and at the same time a reduction of rapsyn by 60% (Figure 7C). In contrast to that, NUP88 downregulation had no effect on MuSK levels, another key player in FADS (20), and no effect on known NUP88 targets, such as CRM1 and NF-κB levels. Similarly, reduced rapsyn levels were observed in C2C12 cells depleted for NUP88 using siRNAs (Figures 7B and C) and shRNAs (Figure 7C). Muscle biopsy of affected individual B.II.2 and immunohistochemistry on paraffin sections furthermore showed weaker staining and irregular distribution of rapsyn in the cytoplasm in comparison to biopsy samples from a control fetus (Figure 7D). Rapsyn is known to not only localize to the plasma membrane, but also to the cytoplasm (34–36). Rapsyn protein levels could not be determined in zebrafish due to a lack of antibodies, but qRT-PCR analyses revealed a reduction of *RAPSN* mRNA by about 20% (Figure 7E).

**Figure 7:**
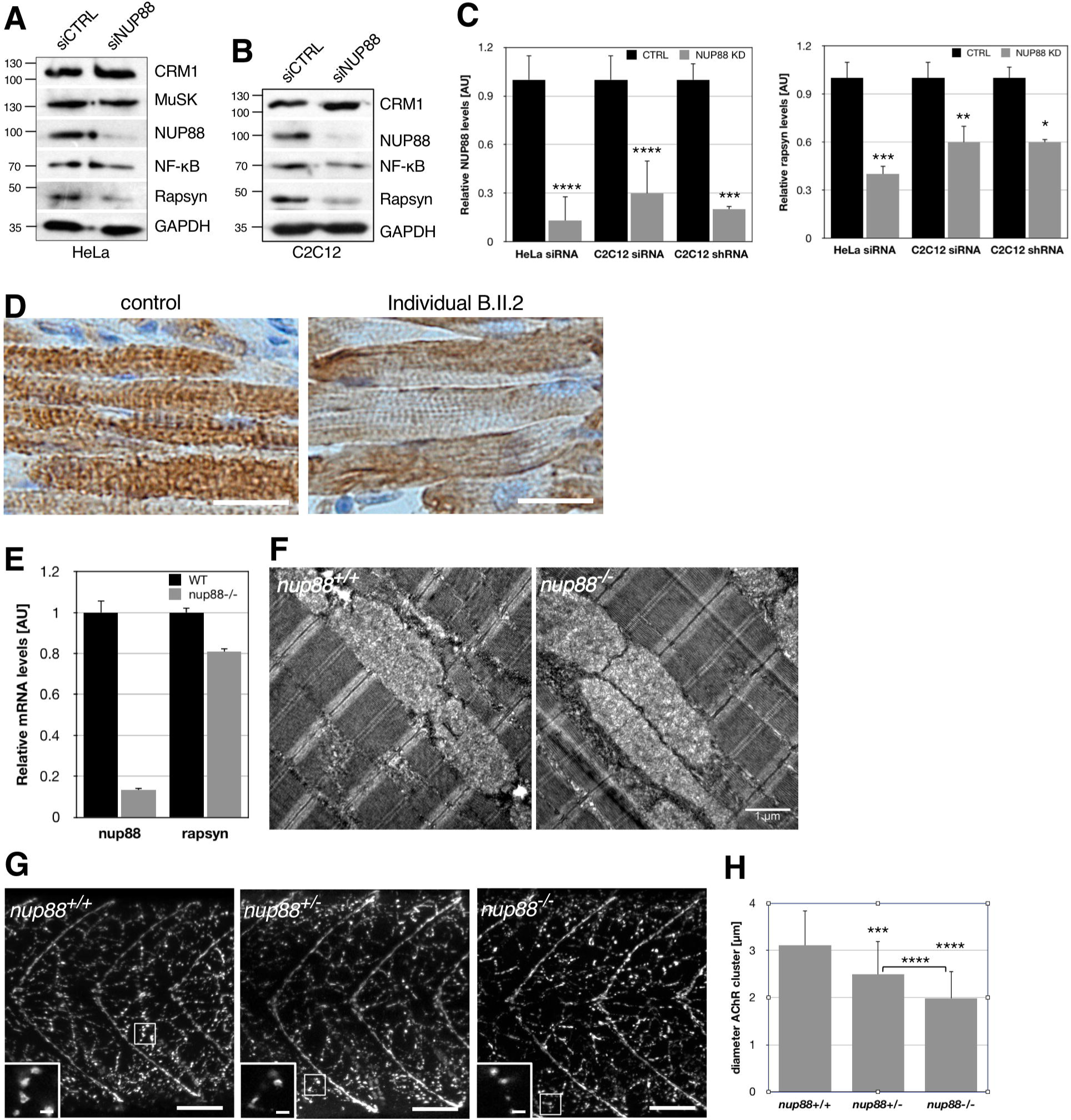
Loss of functional NUP88 affects rapsyn expression as well as AChR clustering. (**A**) HeLa and (**B**) C2C12 cells were treated with the indicated siRNAs for 2 days and cellular lysates were subjected to Western blot analysis using antibodies against NUP88, CRM1, MuSK, NF-kB, and rapsyn. GAPDH was used as loading control. Rapsyn protein levels are reduced in NUP88-depleted cells. Note MuSK has a predicted molecular weight of 97 kDa, but migrates higher (20). (**C**) Quantification of the respective expression levels of NUP88 and rapsyn after transfection of HeLa and C2C12 cells with the indicated siRNAs and shRNA-mediated depletion of NUP88 in C2C12 cells. Blots from three independent experiments for each condition were analyzed. Data present mean ± SEM. P-values ****<0.0001, ***<0.001; **<0.01, *<0.05; t-test, one-tailed. (**D**) Bright-field images of histological muscle sections from individual B.II.2 and a control fetus stained with anti-rapsyn antibodies (brown). Nuclei were visualized by hematoxylin. (**E**) qRT-PCR analysis of nup88 and rapsyn transcripts in wild-type and *nup88^−/−^* embryos. (**F**) Skeletal muscle organization remains unaffected in *nup88^−/−^* zebrafish larvae. Larvae were prepared at 5 dpf for transmission electron microscopy. Skeletal muscle of WT and mutant zebrafish show intact myofibril alignment with their regularly stacked Z-line and clearly identifiable H- and I-zones. Shown are representative longitudinal sections of skeletal muscle from *nup88^+/+^* and *nup88^−/−^* larvae, respectively. Scale bar, 1µm. (**G**) Anterior trunk regions of wild-type (left) and *nup88* heterozygous and homozygous mutant larvae were stained with antibodies against the AChR and secondary Alexa 488 antibodies. AChR cluster size was significantly reduced in *nup88^−/−^* larvae as compared to *nup88^+/+^* and *nup88^+/−^* larvae (higher magnification insets). Insets are taken from the marked area in the respective overview image. Inset for *nup88^−/−^* was rotated by 180°. Scale bars, 50 µm (overview), 5 µm (insets). (**H**) Quantification of the AChR size in *nup88^+/+^* (n=8), *nup88^+/−^* (n=14), and *nup88^−/−^* (n=12) larvae. Per larvae 100 clusters throughout z-stacks of confocal images were manually measured using ImageJ. Data present mean ± SD. P-values ****<0.0001, ***<0.001; t-test, one-tailed.

Consistent with reduced rapsyn levels, we observed impaired AChR clustering in fast-twitch muscle fiber synapses, but not in myoseptal synapses of the 5 dpf zebrafish trunk (Figure 7G). Quantification of the size of individual AChR cluster in WT and mutant zebrafish revealed that the diameter of the AChR clusters was significantly reduced in *nup88^−/−^* larvae as compared to *nup88^+/+^* larvae (Figure 7H). Interestingly, AChR cluster size was also reduced in *nup88^+/−^* larvae, both in comparison to WT and *nup88^−/−^* larvae. This reduced size of AChR clusters in the heterozygotes may account for the observed defects in touch-evoke response (Figure 4H). In accordance with impaired neuromuscular junction formation as a consequence of loss of *nup88*, muscle organization in zebrafish appeared indistinguishable in electron micrographs from *nup88^+/+^* and *nup88^−/−^* larvae (Figure 7F). Similarly, in affected fetus B.II.2 skeletal muscle structure was, based on the autopsy report, intact. Thus, both *in vitro* and *in vivo* evidence support the notion that loss-of-function of NUP88 has a negative effect of on rapsyn, which likely affects AChR clustering and proper formation of neuromuscular junctions.

## Discussion

Here, we have identified biallelic homozygous and compound heterozygous mutations in *NUP88* as cause of fetal akinesia. We demonstrate that the mutations in *NUP88* lead to a loss-of-function phenotype, which coincides with reduced spontaneous motor activity and touch-evoked escape response in zebrafish. Consistent with the fact that the mutations in NUP88 affect different regions of the protein, we observed distinct effects of the mutants on binding to NUP214 and NUP62 in GFP-trap assays, whereas binding to NUP98 appears indistinguishable between wild-type and mutant forms of NUP88 (Figure 6A). This suggests that impaired interaction with partner nucleoporins may contribute, but are unlikely to be causative for NUP88 malfunction in FADS. Our data further suggest that NUP88 malfunction in FADS is at least in part due to dysfunctional rapsyn, a known key player in FADS, and consequently impaired AChR clustering and neuromuscular junction formation. Muscle integrity, in contrast, appears unaffected by a loss of NUP88.

Genetic disruption of *nup88* in zebrafish led to pleiotropic morphological defects, including micrognathia, smaller head and eyes, distortion of the body axis, and aplastic swim bladder (Figures 3-5, Figures S2 and S3), and to impaired locomotor behavior (Figure 4). These phenotypes parallel defects observed in human fetuses affected by FADS, such as reduced fetal movement, micrognathia, joint contractures and lung hypoplasia (for detailed comparison see Table S3). The reduced head and eye size likely originates from increased apoptosis of neuronal cells (Figure 3C) and it will be interesting to identify the affected subset of neurons. *nup88* inactivation affected spontaneous movement of zebrafish from 4 dpf onwards, which resembles the onset of symptoms in the affected fetuses at about week 18 of gestation. The impaired touch-evoked response seen in the mutant zebrafish further matches the absence of reflex response observed in at least some fetuses (Family A, I.2 and B.F., personal communication). Rescue experiments in zebrafish with wild-type or *nup88* mutants revealed that none of the mutant mRNA could restore a wild-type-like phenotype (Figures 5 and S3), indicating that *NUP88* mutations in individuals affected with FADS are loss-of-function variants. The zebrafish model developed here therefore provides a valuable *in vivo* system to further test *nup88* deficiency. This in turn will be key in understanding the role of NUP88 in the etiology of FADS and its function in embryonic development.

Reduced rapsyn levels might partly cause fetal akinesia upon loss of NUP88. Rapsyn is one of the many contributing proteins required for the correct assembly of the AChR and is particularly involved in AChR assembly and localization to the cell membrane (37, 38). We observed reduced rapsyn protein levels in the absence of functional NUP88 in cellular assays using human and mouse cell lines in combination with siRNA and shRNA-mediated depletion of NUP88 (Figures 7). Due to a lack of cell lines derived from affected individuals, we could analyze rapsyn only in histological sections from muscle of one affected fetus, which revealed a weaker staining for and a perturbed intracellular localization of rapsyn (Figure 7D). Consistent with aberrant rapsyn expression, AChR clustering in trunk regions of *nup88^+/−^* and *nup88^−/−^* zebrafish was impaired (Figures 7G and 7H). Rapsyn protein levels could not be determined in zebrafish due to a lack of antibodies, but qRT-PCR analyses revealed a reduction of its mRNA by about 20% (Figure 7E). Our data therefore suggest that absence of functional NUP88 causes fetal akinesia at least in part through misregulation of rapsyn expression. How loss of NUP88 results in reduced rapsyn levels on a mechanistic level remains to be seen. Moreover, this likely is not the only pathway by which NUP88 acts in NMJs, as effects on only one cellular pathway would be indeed (i) very surprising for a nucleoporin *per se*, (ii) irreconcilable with the cranial defects observed in human and zebrafish, and (iii) not in line with the broad central nervous system expression pattern of nup88 in zebrafish. Our data, however, demonstrate (i) that the nup88 spectrum of phenotypes indeed include locomotor defects and that therefore NUP88 deficiencies might result in FADS in humans, and (ii) that human alleles are dysfunctional. We further observed that (iii) NMJ defects correlate to this phenotype. Whether this is the primary cause of akinesia is impossible to determine given the pleiotropic effects of nup88 deficiency, but at least they are sufficient to explain the phenomenon. Further mechanistic details will be subject for future studies.

## Methods

### DNA sequencing Family A

Exome sequencing in one affected baby (A.II.7) was performed at the Clinical Exome Sequencing (CES) at University of California, Los Angeles and the sequencing report is on hand. In total 22,843 DNA variants were identified, including 21,625 single nucleotide substitutions and 1,218 small deletions/insertions (1-10 bp): the data were consistent with a high quality genomic sequence and fall within normal human genomic variation quality parameters. Estimated from these data, about 93% of the exome was reliably sequenced with at least 10x coverage. In total, 5 homozygous and 329 rare heterozygous protein-altering variants of uncertain clinical significance were identified across 313 genes. A rare autosomal-recessive model of inheritance with homozygous causative mutations due to consanguinity of the parents was assumed. After applying appropriate filters, a novel homozygous variant c.1300G>T, p.D434Y in the *NUP88* gene [NM_002532.5] was identified in the subject’s DNA. This variant had not been previously observed in the general population and was predicted to be deleterious/probably damaging by three *in silico* prediction algorithms (Table S1). Additionally, the homozygous mutation c.1300G>T, p. D434Y in *NUP88* was confirmed in a second affected fetus A.II.5 by Sanger sequencing.

### DNA sequencing Family B

Exome sequencing was performed on DNA from the proband B.II.2, both healthy sisters and both parents from the European family, as outlined previously (39). Exome enrichment was performed on DNA using an Ampliseq Whole Exome kit, (Thermofisher Scientific). Briefly, a total of 100 ng of DNA was amplified in 12 separate PCR pools, each containing ~25,000 primer pairs. After amplification, the individual reactions were pooled and digested to degrade the PCR primers. Next, barcoded sequencing adaptors were ligated and the library was purified using AMPure beads (Beckman Coulter), amplified and purified again and analyzed on a 2100 Bioanalyzer (Agilent Technologies). Libraries were diluted to 18-26 pM and attached to Ion Sphere Particles (ISPs) using an Ion Proton Template 200 V3 kit and sequenced on a P1 sequencing chip for 520 flows on an Ion Proton sequencer (Ion Sequencing 200 kit V3). Two samples were pooled and sequenced on a single chip. Following sequencing, reads were trimmed to remove low quality bases from the 3’ end and mapped to the human genome reference sequence (HG19) using tmap (Torrent Suite 4.2). Variant calling was performed using the Torrent Variant Caller with custom settings optimized for whole exomes and the data was annotated using ion Reporter 4.0. Variants were filtered with ANNOVAR (40) against ENCODE GENECODE v.19, 1000genomes (threshold >0.5%), dbSNP138 common databases and against a list of in-house common variants. Genes with variants that fitted an autosomal recessive inheritance pattern and co-segregated with disease in the family were prioritized.

*NUP88* was the only candidate gene that harbored variants compatible with an autosomal recessive inheritance pattern in this family (Figure S1B). Individual B.II.2 was compound heterozygous for an in-frame 3 bp deletion in exon 14 (c.1899_1901del, p.E634del) and a nonsense mutation c.1525C>T, p.R509* in exon 11. Mutation c.1899_1901del mutation was inherited on the maternal allele and the c.1525C>T paternally. Healthy sister B.II.1 is heterozygous for the maternal mutation, healthy sister B.II.3 is a heterozygous carrier of the paternal mutation (Figure 1A, pedigree Family B).

### Structure modelling

Human NUP88 protein sequence Q99567 (UniProt database) was used for structural investigation and modelling using an N-terminal and C-terminal part, according to predictions. Database search comparisons using HHPRed (41–45) revealed as best hits for the N-terminal domain (NTD): Nuclear pore complex proteins Nup82 from *S. cereviseae* and *Chaetomium thermophilum* (PDBid: 3pbp A, 3tkn_A and 5cww_B, respectively) and for the C-terminal domain (CTD) Nup57 from *Chaetomium thermophilum* (PDBid: 5cws E) and Nup54 from Homo sapiens (PDBid: 5ijn F). The high similarity to yeast and *Chaetomium* Nup82, the structures of which have been solved in part (46–48) strongly indicate that the NTD also forms a ß-propeller and predictions suggest that the CTD is in an all helical arrangement (PsiPred and JPred4) (49, 50). In order to gain insight into the putative fold and the location of the disease-related residues identified in hsNUP88, four servers were used for structure prediction. Phyre2, Robetta, I-Tasser, RaptorX (51–57) were supplied with either full length sequence or only the NTD (residues 1-495) or CTD (residues 496-741) in case of residue limitations or implausible models resulting from full length submission. All modelling was performed using standard settings. All figures were generated using Pymol.

### Zebrafish husbandry

Zebrafish (*Danio rerio*) were raised and bred at 28°C on a 14 hr/10 hr light/dark cycle. Embryos and larvae were raised in egg water (0.3 g/l Instant Ocean Salt, 75 mg/l CaSO_4_; 1 mg/l Methylene Blue). The line carrying *nup88*^sa2206^ allele was obtained from Zebrafish Mutation Project, Knockouts for Disease Models (http://www.sanger.ac.uk/sanger/Zebrafish_Zmpgene/ENSDARG00000003235#sa2206). Heterozygous fish for *nup88* were out crossed for four generations with wild-type fish before analysis. All animal experiments were performed in accordance with the rules of the State of Belgium (protocol approval number: CEBEA-IBMM-2012:65).

### RNA expression constructs

Capped messenger RNA was synthesized using the mMESSAGE mMACHINE kit (Ambion). The following expression plasmids were generated and used in this study: the full-length zebrafish *nup88* ORF was cloned from 48 hpf cDNA and recombined into BamHI-XhoI digested pCS2 using In-fusion cloning (Takara). *nup88* mutants corresponding to the sequences identified in human fetal akinesia cases were generated by site-directed mutagenesis using QuikChange Lightning Site-Directed Mutagenesis Kit (Agilent Technologies). All primer sequences are listed in Table S2. mRNAs (300 pg) were injected at the one-cell stage.

### Genotyping and RT-PCR

nup88^sa2206^ genotyping was performed by RFLP assay using MseI restriction of a 150 bp PCR product. Real-time PCR was done at various stages of embryonic and larval development (4-cell to 5 dpf). Primer sequences are listed in Table S2. Staging of embryos was performed according to (58).

### Alcian blue staining

Alcian Blue staining and histology were performed as described elsewhere (59). All images were acquired using an Olympus SZX16 stereomicroscope and an Olympus XC50 camera using the imaging software Cell* after embryo anesthesia with a low dose of tricaine.

### Apoptosis assay using acridine orange in live embryos

Live embryos at stages 24 hpf, 36 hpf, 48 hpf were dechorionated and immersed in egg water containing 5 μg/ml acridine orange. They were incubated at 28.5°C for 15 minutes in the dark and then thoroughly washed with egg water. Embryos were mounted in low-melting agarose for positioning and immediately imaged using an Axio Observer Z1-1 microscope. Images were processed using Zeiss Zen^TM^ software.

### Behavioral assays in zebrafish embryos and larvae

Spontaneous tail coiling of 22-24 hpf embryos, placed in the grooves of an agarose chamber, was recorded for 2 minutes at 30 frames per second. Coiling events were scored manually. To analyze spontaneous locomotion, 4 dpf larvae were placed into 96-well plates (one larva per well). Their behavior was recorded at 30 frames per second for 5 minutes and quantified using EthoVision XT 8.5 software (Noldus).

Touch-induced escape responses were analyzed in 4 dpf head-restrained larvae. Larvea were first embedded in 2% low melting point agarose, and then the agarose surrounding their tail was removed with a blade. Escape behavior was induced by touching the tail of larvae with a plastic pipette tip and recorded at 300 frames per second. Duration of behavioral responses was quantified with ImageJ.

### qRT-PCR on 5 dpf zebrafish larvae

Total RNA of 5 dpf nup88 mutant and WT zebrafish larvae was extracted using TRIzol reagent (Invitrogen, Carlsbad, CA, USA) according to the manufacturer’s protocol. Genomic DNA was eliminated prior to reverse transcription using PrimeScript RT Reagent Kit with gDNA Eraser (Perfect Real Time; RR047A, Takara, Dalian, China). Total RNA (5⍰μg) was next reverse-transcribed in 20⍰μl final volume according to manufacturer’s protocol. Primers for target genes (Supplementary Table 2) were designed with Realtime PCR Tool by Integrated DNA Technologies, Inc (https://eu.idtdna.com/scitools/Applications/RealTimePCR), and sequences were submitted to BLAST (http://blast.ncbi.nlm.nih.gov/Blast.cgi).

Quantitative PCR was then performed using SYBR^®^ Premix Ex Taq TM II (Tli RNaseH Plus), Bulk kit (RR820L; Takara, Dalian, China). 1 μl of cDNA was used in each 20⍰μl-PCR well with 300 nM primers pair final concentration. Each sample was assayed in triplicate and samples not reverse-transcribed were used as negative controls. Eight housekeeping genes were tested according to (60) and the most stable combination was determined using qbase+^TM^ qPCR analysis software (https://www.qbaseplus.com). bactin2, gapdh and ybx1 were the best combination of housekeeping genes at this stage of larval development and used for normalization.

Differences in amplification curves between the target genes and housekeeping genes were identified by comparing standard curve slopes. Real-Time PCR was performed using Applied Biosystems StepOnePlus™ Real-Time PCR System and PCR analyses were performed with qbase+ system software.

### Zebrafish immunostaining

For whole-mount immunostaining, zebrafish embryos were fixed in 4% PFA for 3 hrs at room temperature (RT), washed and permeabilized with proteinase K (40µg/ml) at 37°C for 1 h. For blocking, embryos were gently shaken in PBS containing 10% of normal goat serum, 1% DMSO and 0.8% Triton X-100 for 1 h and then, incubated with the primary anti-acetylcholine receptor antibody (mouse mAb35 (DSHB, Hybridoma Bank; 1:100) overnight at 4°C. After washing, embryos were incubated with the secondary antibody (anti-mouse IgG Alexa Fluor 594, 1:1000) for 2 h at RT and washed six times for 15 min in PBST prior to mounting and confocal imaging (Zeiss LSM710). Size of AChR cluster were determined using ImageJ.

### Transmission electron microscopy of zebrafish skeletal muscle

Skeletal muscle of 5 dpf nup88^+/+^ and nup88^−/−^ zebrafish embryos were analysed by transmission electron microscopy on longitudinal and transversal ultrathin sections. Briefly, embryos were fixed in 2.5% glutaraldehyde and 0.1 M sodium cacodylate buffer, pH 7.4 overnight at 4°C and post-fixed in 1% osmium tetroxide, 1.5% ferrocyanide in 0.15 M cacodylate buffer for 1 hour at RT. After serial dehydration in increasing ethanol concentrations, samples were embedded in agar 100 (Agar Scientific Ltd., UK) and left to polymerize for 2 days at 60⍰°C. Ultrathin sections (80 nm thick) were collected using a Leica EM UC6 ultramicrotome and stained with uranyl acetate and lead citrate. Images were recorded on a Tecnai10 electron microscope (FEI) equipped with an Olympus VELETA camera and processed with using AnalySIS software.

### Immunohistochemistry-paraffin (IHC-P) labelling of fetal muscle sections

Brain and skeletal muscle paraffin-embedded blocks were obtained from autopsy of individual B.II.2. Written consent form for use of paraffin samples for functional analysis was signed by the parents. Control tissues of a fetus of the same age without neuromuscular disorder were used for IHC-P. Blocks were cut with a microtome and 5 µm sections were mounted. Sections were deparaffinized and rehydrated. Heat-induced epitope retrieval was performed using Tris/EDTA, pH 9.0 for all tissues when using antibodies against rapsyn and with sodium citrate, pH 6.0 for anti-nucleoporin antibody mAb414. 0.3% H_2_O_2_ for 20 minutes was used to block samples endogenous peroxidase followed by washes with PBS. Tissues were permeabilized using PBS containing 0.5% Triton X-100 for 5 minutes at RT and subsequently blocked with PBS containing 1% BSA for 30 minutes. Primary antibodies were diluted in blocking buffer and incubated overnight at 4°C in a humidification chamber. Samples were washed in PBS for 10 minutes. Biotinylated goat anti-rabbit-Ig (1:400; DAKO-E0432) or biotinylated goat anti-mouse (1:400; DAKO-E0433) were added for 30 minutes.

After washing in PBS, samples were incubated 30 minutes with streptavidin-HRP (DAKO-P0397) diluted 1:300 in PBS and thoroughly washed with PBS. DAKO Liquid DAB+ Substrate Chromogen System K3468 (1ml substrate + 1 drop DAB) was added on samples and left until a brown coloring appeared (maximum 6 minutes). After several PBS washes, the samples were counterstained with hematoxylin for 15 seconds, washed thoroughly under running tap water to remove excess staining agent and mounted for subsequent observations with Aquatex. Images were acquired with an Olympus BX41 microscope and were processed using Olympus cellSense^TM^ software.

### Plasmids for studies in human cell lines

For all constructs, human *NUP88* was amplified by PCR. All constructs were verified by DNA sequencing. GFP-NUP88 was produced as described previously (2). FLAG-NUP88 was cloned into *KpnI/XbaI* cut *pFLAG-CMV2* (Sigma-Aldrich). GFP-Nup88 and FLAG-NUP88 mutants were generated by site-directed mutagenesis using the QuikChange Lightning site-directed mutagenesis kit (Agilent Technologies) following the manufacturer’s instructions. Primers are listed in Table S2.

### Antibodies

The following polyclonal antibodies were used in this study for Western blotting (WB), immunofluorescence (IF) and immunohistochemistry (IHC): rabbit anti-Nup88 (kind gift of Dr. Ulrike Kutay; IF 1:1000), rabbit anti-NUP214 (Abcam, ab70497; WB 1:5000, IF 1:500), rabbit anti-lamin A (Sigma-Aldrich, L1293; WB 1:500), rabbit anti-rapsyn (Novus Biologicals, NBP1-85537; WB 1:500; IHC 1:200), rabbit anti-MuSK (ThermoScientific, PA5-14705; WB 1:1000), rabbit anti-GAPDH (Cell Signaling, 2118; WB 1:10.000), rabbit anti-actin (Sigma-Aldrich, A2066; WB 1:1000), rabbit anti-Nesprin1 (Sigma-Aldrich HPA019113; IF 1:250), rabbit anti-Nesprin2 (a kind gift of Dr. Iakowos Karakesisoglou; IF 1:50), rabbit anti-Sun1 (kind gift of Dr. Ulrike Kutay; IF 1:1000), rabbit anti-Sun2 (Sigma-Aldrich, HPA001209; IF 1:200), rabbit anti-emerin (Abcam, ab153718; IF 1:500)

The following monoclonal antibodies were used in this study: mouse anti-Nup88 (BD Transduction Laboratories, 611896; IF 1:500), mouse mAb414 (Covance MMS-120R; IF 1:2000; IHC 1:100), mouse anti-NUP62 (Clone 53, BD Transduction Laboratories, 610497; WB 1: 3000, IF1:500), mouse anti-lamin A/C (Abcam, ab8984; WB 1:300, IF 1:30), mouse anti-FLAG (Sigma-Aldrich F-3165; IF: 1:200), rat anti-NUP98 (Sigma-Aldrich, N1038; WB 1:2000, IF 1:1000), mouse anti-AChR (mAb35, DSHB, Hybridoma Bank; IF 1:100), rat anti-GFP (3H9, Chromotek; WB 1:1000).

Secondary antibodies for immunofluorescence were the corresponding goat anti-mouse-IgG Alexa 568 (1:1000; Invitrogen), goat anti-rabbit IgG Alexa 568 (1:1000; Invitrogen). Secondary antibodies were either alkaline phosphatase coupled antibodies from Sigma/Aldrich and used at 1:20.000 or HRP coupled antibodies from Cell Signaling Technology at 1:8000.

### Cell culture and transfections

All experiments were conducted in HeLa cells (provided by Robert D. Goldman, Feinberg School of Medicine, Northwestern University, Chicago, USA) grown in Dulbecco’s modified Eagle’s medium (DMEM), supplemented with 10% fetal bovine serum (FBS) plus penicillin and streptomycin. Cells were transfected with plasmids using Turbofect transfection reagent (Thermo Scientific Fermentas, St. Leon-Rot, Germany), Lipofectamine 2000 (Life Technologies Invitrogen, Gent, Belgium) or jetPRIME (Polyplus, Illkirch, France) and with siRNAs using Lipofectamine RNAiMAX (Life Technologies Invitrogen) following the instructions of the manufacturer. Smart-pool small interfering RNAs were obtained from Dharmacon (GE Healthcare Europe, Diegem, Belgium): human NUP88 (L-017547-01-0005), mouse NUP88 (L-054949-01-0005), and non-targeting siRNAs (D-001810-10).

### Immunofluorescence microscopy of HeLa cells

HeLa cells were grown on glass coverslips, transfected, fixed in 4% PFA in PBS for 5 min, permeabilized with 0.5% Triton-X-100 in PBS for 5 min and then fixed again. Blocking was performed with 2% BSA/0.1% Triton-X-100 in PBS for 30 min at RT. Primary antibodies were incubated at 4°C over-night in a humidified chamber. Secondary antibodies were incubated 1 hour at RT in the dark. Excess antibodies after primary and secondary antibody staining were removed by three washing steps using 0.1% Triton-X-100 in PBS for 5 min. Cells were imaged using a Zeiss LSM 710 (Zeiss, Oberkochen, Germany) confocal laser scanning microscope with Zeiss Plan-Apochromat 63x/1.4 oil objective. Images were acquired using the microscope system software and processed using Image J and Adobe Photoshop (Adobe Systems, Mountain View, CA).

### GFP-trap assay

HeLa cells, grown in a 10 cm dish, were transfected with GFP and GFP-NUP88 wild-type and mutant variants, respectively. Cells were grown for 48 hours at 37°C in a humidified atmosphere with 5% CO_2_. To harvest cells, growth medium was aspirated off, 1 ml of ice-cold PBS was added to cells and cells were scraped from dish. The cells were transferred to a pre-cooled tube, spinned at 500 g for 3 min at 4°C and the supernatant was discarded. The cell pellet was washed twice with ice-cold PBS. Pellets were subsequently lysed with 200 μl of ice-cold lysis buffer (10 mM Tris/Cl, pH 7.5, 150 mM NaCl, 0.5 mM EDTA, 0.5 % NP-40, protease-phosphatase inhibitor) using a syringe before incubating on ice for 30 minutes. The tubes were centrifuged for 15 minutes at 16.000 g at 4°C and the supernatant was transferred into a new reaction tube.

Bradford assay was used to determine the protein concentration of the lysates and 300 μg of protein lysate adjusted to 500 μl in dilution buffer (10 mM Tris/Cl, pH 7.5, 150 mM NaCl, 0.5 mM EDTA, protease-phosphatase inhibitor) was added to 25 μl of GFP-Trap^®^_MA beads (ChromoTek, Planegg-Martinsried, Germany). Beads were prewashed twice with dilution buffer. The beads and the lysates were incubated 1hour at 4°C on an end-to-end rotor. The magnetic beads were then washed three times in dilution buffer containing 150 mM, 250 mM and 500 mM NaCl following the addition of 20 μl of 2x SDS-sample buffer (120 mM Tris/Cl, pH 6.8, 20% glycerol, 4% SDS, 0.04% bromophenol blue, 10% β-mercaptoethanol) and boiling at 95 °C for 10 minutes. The eluates were subsequently loaded on to 7% polyacrylamide gels and Western blot was carried out (see below).

## Western blotting

HeLa and C2C12 (provided by Vincent Mouly, The Pitié-Salpêtrière Hospital, Institute of Myology, Paris, France) cells were lysed in lysis buffer (50 mM Tris-HCl, pH 7.8, 150 mM NaCl, 1% Nonidet-P40 and protease inhibitor cocktail tablets (Roche, Basel Switzerland)). 20 µg of protein were loaded and separated by sodium dodecyl sulfate-polyacrylamide (5% or 7%) gel electrophoresis (SDS-PAGE). The proteins were transferred onto a PVDF membrane (Immobilon-P, Millipore) and the membranes were blocked with TBS containing 0.1% Tween 20 and 5% non-fat dry milk for 1 hour. The membranes were then incubated for 1 hour in blocking solution containing a primary antibody followed by washing 3x in TBS containing 0.1% Tween 20 and 5% non-fat dry. The membranes were next incubated with secondary antibodies for 1 hour, washed 3x in TBS and developed. X-ray films were scanned and processed using ImageJ.

## Author contributions

BF, BV, EB, AF, GR, KH, RJ, MH, and LP wrote the manuscript. BF, BV, and KH designed and supervised the study. EB, PC, AF, RJ, SK, MH, AD, VM, MV, LS, NM, LP, GR, ML, SP, CS, SJ, ER, JVD, ML, RHK, RF, NL, KH, BV, and BF acquired, analyzed and interpreted the data.

## Acknowledgements

We are grateful to the families of the affected babies for participating with this work, clinicians and genetic counsellors for the sample collection and Clinical Exome Sequencing (CES) at UCLA for sharing exome sequencing results. We thank Dr. Robert Bryson-Richardson (Monash University, Australia) for advice concerning zebrafish muscle sections. We are grateful to Ulrike Kutay (ETH Zürich, Switzerland), Iakowos Karakesisoglou (Durham University, UK), and Shawn Ferguson (Yale School of Medicine, New Haven, USA) for sharing reagents. Confocal and TEM images were acquired at the CMMI, which is supported by the European Regional Development Fund (ERDF).

This work was supported by grants from the Fédération Wallonie-Bruxelles (ARC 4.110.F.000092F), the Fonds Brachet and the Fonds Van Buuren to BF. and by a fellowship of Fonds Hoguet and Fonds Brachet to EB. Work in the B.V. laboratory is supported by the Fonds De La Recherche Scientifique (FNRS) (MIS F.4543.15), an ARC grant, the Fondation ULB, the Queen Elisabeth Medical Foundation for Neurosciences (Q.E.M.F.,) and the Fonds de la Recherche Scientifique - FNRS for the FRFS-WELBIO (CR-2017S-05). P.C. is a Postdoctoral Researcher from the FNRS. GR is supported by an Australian National Health and Medical Research Council (NHMRC) Career Development Fellowship (APP1122952), NGL by NHMRC Principal Research Fellowship (APP11117510). This work was funded by a NHMRC Project Grant (APP1080587) and the AFM (15734). S.P and R.H.K were supported by a grant from the DFG (SFB860).

